# Single-nucleotide-resolution genomic maps of *O*^6^-methylguanine from the glioblastoma drug temozolomide

**DOI:** 10.1101/2023.12.12.571283

**Authors:** Jasmina Büchel, Cécile Mingard, Vakil Takhaveev, Patricia B. Reinert, Giulia Keller, Tom Kloter, Sabrina M. Huber, Maureen McKeague, Shana J. Sturla

**Affiliations:** Department of Health Sciences and Technology, ETH Zurich, Zurich 8092, Switzerland; Department of Chemistry, McGill University, Montreal H3A 0B8, Canada

## Abstract

Temozolomide kills cancer cells by forming *O*^6^-methylguanine (*O*^6^-MeG), which leads to apoptosis due to mismatch-repair overload. However, *O*^6^-MeG repair by *O*^6^-methylguanine-DNA methyltransferase (MGMT) contributes to drug resistance. Characterizing genomic profiles of *O*^6^-MeG could elucidate how *O*^6^-MeG accumulation is influenced by repair, but there are no methods to map genomic locations of *O*^6^-MeG. Here, we developed an immunoprecipitation- and polymerase-stalling-based method, termed *O*^6^-MeG-seq, to locate *O*^6^-MeG across the whole genome at single-nucleotide resolution. We analyzed *O*^6^-MeG formation and repair with regards to sequence contexts and functional genomic regions in glioblastoma-derived cell lines and evaluated the impact of MGMT. *O*^6^-MeG signatures were highly similar to mutational signatures from patients previously treated with temozolomide. Furthermore, MGMT did not preferentially repair *O*^6^-MeG with respect to sequence context, chromatin state or gene expression level, however, may protect oncogenes from mutations. Finally, we found an MGMT-independent strand bias in *O*^6^-MeG accumulation in highly expressed genes, suggesting an additional transcription-associated contribution to its repair. These data provide high resolution insight on how *O*^6^-MeG formation and repair is impacted by genome structure and regulation. Further, *O*^6^-MeG-seq is expected to enable future studies of DNA modification signatures as diagnostic markers for addressing drug resistance and preventing secondary cancers.

**GRAPHICAL ABSTRACT:** 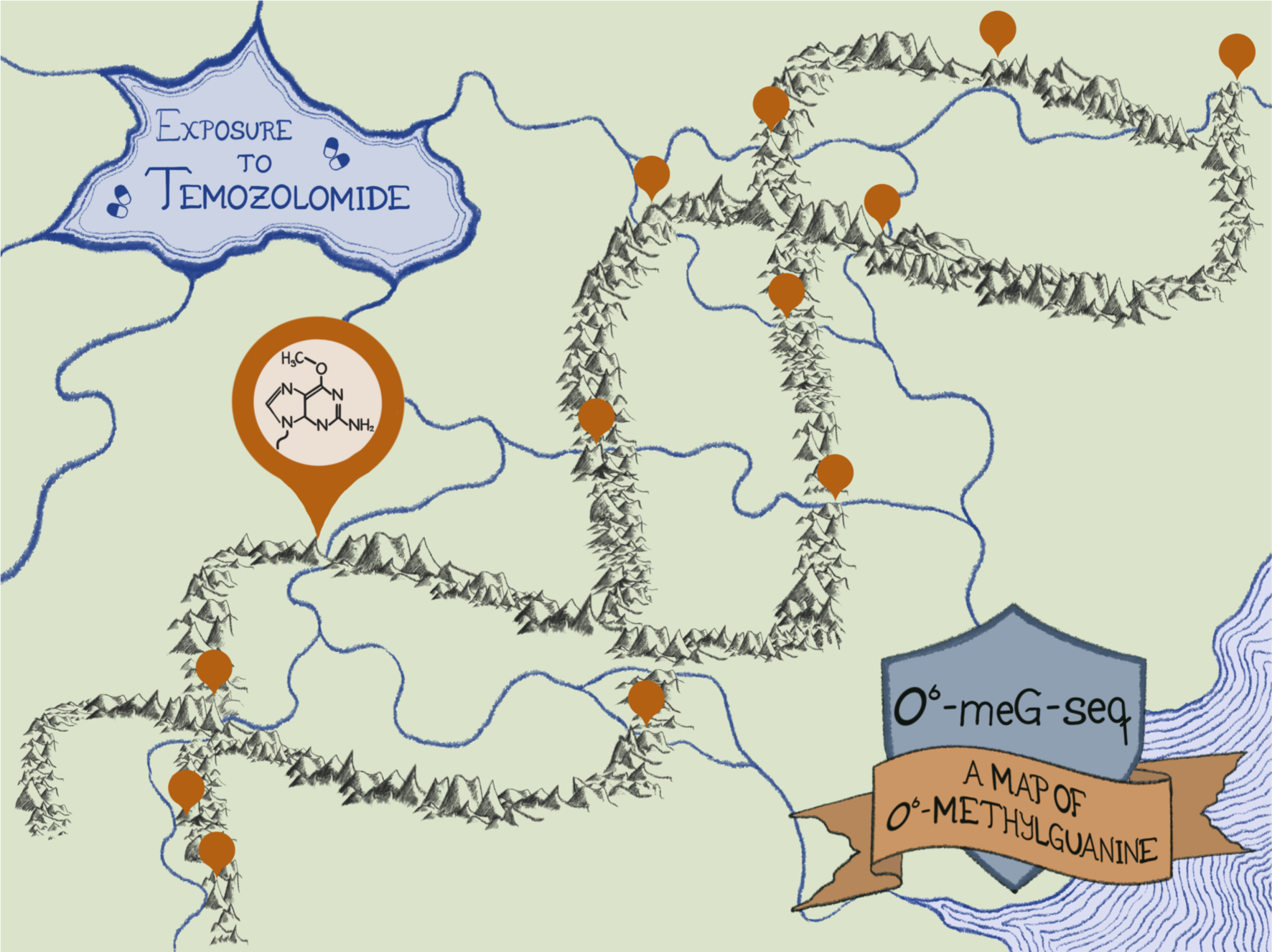

## INTRODUCTION

Glioblastoma is a malignant brain tumor affecting 2-5 people per 100’000 every year with a median survival of 12-15 months.(1,2) The current standard of care is surgery and radiotherapy combined with temozolomide (TMZ), a DNA alkylating agent that effectively crosses the blood-brain barrier.(3) However, over 50% of patients treated with TMZ do not respond due to enzymatic repair of TMZ-induced alkylation in target cells.(1,4,5) There is little knowledge regarding the landscape of TMZ-induced alkylation or TMZ-resistance-associated repair throughout structural or functional genome features because there are no available genomic maps of TMZ-induced DNA alkylation, which define the occurrence of chemical modification at each base in the human genome.

TMZ functions by inducing DNA alkylation adducts, such as *O*^6^-methylguanine (*O*^6^-MeG)(6), causing mismatches upon replication due to mispairings with thymine instead of cytosine. Mismatch repair (MMR) removes incorrectly inserted nucleotides opposite *O*^6^-MeG, but since it does not remove *O*^6^-MeG itself, mispairing continues and MMR repeatedly creates gaps in the newly synthesized strand.(7) Thus, persistent *O*^6^-MeG leads to a futile cycle of MMR, which triggers apoptosis.(8) In contrast, *O*^6^-MeG can be directly repaired by *O*^6^-methylguanine-DNA methyltransferase (MGMT, also known as *O*^6^-alkylguanine-DNA alkyltransferase AGT).(8,9) MGMT transfers the methyl group from DNA to an active site cysteine residue (9), directly leaving behind a repaired guanine. As a result of repair, MGMT-expressing tumor cells are likely to be resistant to chemotherapy with TMZ.(1,10) While profiling MGMT promotor methylation status is a widely used diagnostic for anticipating TMZ sensitivity or resistance, drug resistance remains a major limitation for glioblastoma therapy.

Patients with epigenetically methylated *MGMT* promoters, and therefore low MGMT expression, are more likely to survive glioblastoma (10) but may be at increased risk for secondary cancers due to the mutagenic effects of *O*^6^-MeG.(11,12) One of the mutational signatures annotated in the catalogue of somatic mutations in cancer (COSMIC), namely single base substitution (SBS) 11, has been identified in genomes of secondary cancers from patients previously treated with TMZ.(11,12) SBS 11 is strongly characterized by C-to-T mutations hypothesized to arise from TMZ-induced *O*^6^-MeG. However, how DNA modification signatures, specifically *O*^6^-MeG signatures, in TMZ-exposed cells relate to SBS 11 is not known since no such TMZ-induced DNA modification signatures have been reported.

DNA modification signatures involve patterns of modified DNA in certain sequence contexts as putative precursors of mutational signatures. DNA modification signatures as well as mutational signatures not only reflect how chemicals modify DNA but also how repair pathways influence the lesion context.(13) Exploring these relationships has become possible recently as novel methods have emerged for sequencing different types of DNA modifications (14) ranging from small modifications like 8-oxoguanine (8-oxoG) (14–16) to alkylation and drug-induced DNA adducts.(17,18) A common mapping strategy is to utilize anti-adduct antibodies to enrich for DNA fragments containing a specific DNA adduct, followed by marking the exact position of the DNA adduct using a stalled high-fidelity polymerase.(18) By this strategy, cisplatin DNA crosslinks were observed to form uniformly across the genome but their steady state accumulation was driven by repair efficiency.(18,19) While this approach has proven to be highly versatile and it has been adapted to map various adducts such as cisplatin crosslinks (18), UV pyrimidine dimers (20,21), and benzo(a)pyrene adducts (17), it is limited to bulky DNA adducts that readily stall DNA polymerases. A limitation to using this common strategy to map smaller modifications, such as *O*^6^-MeG arising from TMZ, concerns their proficient bypass by many polymerases, including the high-fidelity DNA polymerase Q5 used in previous research.(17–22)

In this work, we created the first genome-wide map of *O*^6^-MeG in a human glioblastoma cell line and characterized how *O*^6^-MeG distribution in the genome, as well as drug sensitivity, is impacted by MGMT repair. First, we screened high-fidelity DNA polymerases for their capacity to stall at small modifications and found the high-fidelity DNA polymerase Platinum SuperFi II to stall at *O*^6^-MeG. This observation allowed us to establish a new method termed *O*^6^-MeG-seq and use it to precisely locate *O*^6^-MeG in wild type LN229 glioblastoma cells (MGMT deficient) exposed to TMZ. To determine the potential impact of MGMT on *O*^6^-MeG genome distribution, we also applied *O*^6^-MeG-seq to characterize TMZ-exposed LN-229 cells transfected with an *MGMT* harboring plasmid. We extracted *O*^6^-MeG signatures from trinucleotide patterns of both cell lines upon TMZ exposure and compared them to known SBS signatures. Additionally, we investigated genome-wide patterns of *O*^6^-MeG distribution and compared them to chromatin accessibility. Finally, we determined how MGMT influenced the preferred accumulation of O6-MeG in gene bodies.

## MATERIAL AND METHODS

### LN-229 cell characterization and TMZ exposure

#### Reagents

Cells were cultured in GlutaMAX™ DMEM (Gibco™, purchased from Thermo Scientific™, 31966047) supplemented with 10% FBS (Gibco, purchased from Thermo Scientific™, 10270106) and 1% Penicillin-Streptomycin (Gibco™, purchased from Thermo Scientific™, 15140122) at 5% CO_2_ and 37 °C. For passaging, cells were detached with Trypsin-EDTA (0.25%), phenol red (Gibco™, purchased at Thermo Scientific™, 25200-056). DNA was extracted using the QIAamp® DNA Mini Kit (Qiagen, 51304). Cell viability was assessed using CellTiter-Glo® Luminescent Cell Viability Assay (Promega, G7571). cOmplete™ EDTA-free Protease Inhibitor Cocktail (Roche, purchased from Sigma, 4693159001), Pierce™ BCA Protein Assay Kits (Thermo Scientific™, 23225), Trans-Blot Turbo RTA Mini 0.2 µm PVDF Transfer Kit (BioRad, 1704272) and Pierce™ enhanced chemiluminescence (ECL) Western Blotting Substrate (Thermo Scientific™, 32109) were used for western blotting in addition to the following antibodies: anti-MGMT mouse antibody (Santa Cruz Biotechnology, SC-56157), anti-Actin rabbit antibody (Sigma A2066), Goat anti-mouse horse radish peroxidase (HRP) antibody (Abcam, ab6789), Goat anti-Rabbit IgG (H+L) Highly Cross-Adsorbed Secondary Antibody, Alexa Fluor™ 488 (Invitrogen™, purchased from Thermo Fisher Scientific, A11034). Fluorescent and chemiluminescent images were taken using a ChemiDoc™ MP Imaging System (Bio-Rad Laboratories, Inc.). Luminescence was measured with a Tecan Infinite 200 PRO^®^ (Tecan Trading, Ltd.). DNA concentration was measured using a Quantus™ Fluorometer (Promega, E6150) with QuantiFluor® ONE dsDNA dye (Promega, E4870). Temozolomide was purchased from Sigma-Aldrich (T2577).

#### Biological Resources

LN-229 cells stably transfected with negative control plasmid (WT) or plasmid containing the MGMT (+MGMT) were provided by Prof. Michael Weller, Zürich University Hospital. Cells were authenticated by Microsynth AG and tested for mycoplasma contamination.

#### Cell viability

The sensitivity of LN-229 WT and LN-229 +MGMT cells to TMZ was tested by measuring intracellular ATP content. LN-229 cells, WT and +MGMT, were seeded in technical triplicates in 96-well plates (8000 cells per well for 24 h, 2000 cells per well for 72 h and 1000 cells per well for 144 h incubation after TMZ exposure). The day after, cells were exposed to increasing TMZ concentration (50 µM to 1 mM) or 1% DMSO solvent control in normal growth medium. Additionally, cells were repeatedly exposed to TMZ by replacing the medium with fresh TMZ-containing medium after 4 and 8 h. For the 144 h incubation, the medium was replaced after 72 h to avoid starvation. In all cases, viability was measured 24, 72, and 144 h after exposure with CellTiter-Glo® Luminescent Cell Viability Assay per manufacturer’s instructions and luminescence was measured. Data was normalized to the solvent control and, for visualization, fitted with LOWESS (Locally Weighted Scatterplot Smoothing) with default settings in statsmodels (version 0.12.0).

#### Western Blot

MGMT expression in LN-229 cells +/- MGMT was assessed by western blotting. LN-229 cells were seeded on 10 cm dishes (1 x 10^6^ cells per dish). When reaching 80% confluency, cells were harvested by first washing them three times with ice-cold PBS and then by adding 100 µL of lysis buffer (final concentrations: 1 mM PMSF, 1 mM Na_3_VO_4_ 10 mM NaF, and 1X EDTA-free Protease Inhibitor Cocktail in RIPA buffer). Cells were scraped and collected in a tube on ice, sonicated twice every 15 minutes. The cell lysate was centrifuged at 16’900 g (4 °C) and the supernatant was collected. Protein quantification was then performed using the Pierce BCA Protein Assay Kit according to the manufacturer’s instructions. Protein samples (33 µg) were mixed with 5X LDS loading buffer (8% SDS, 0.5 M dithiothreitol, 50% glycerol, 0.25 M Tris-HCL pH 6.8, bromophenol blue 0.05 %) to achieve a sample volume of 20 µL. Samples were denatured at 90 °C for 10 minutes. Samples were run on a 4-12% Bis-Tris SDS Protein gel with 1X MES SDS running buffer, 3 µL of PM2610 protein ladder was also loaded. The run was started at 70 V for 30 minutes, then the voltage was increased to 100 V for 2-3 h. Separated proteins were transferred to polyvinylidene fluoride membranes using the Trans-Blot Turbo RTA Midi PVDF Transfer Kit. The membrane was blocked with 5% milk powder in TBS-T (0.05% Tween, 20 mM Tris-HCL, 150 mM NaCl, pH 7.5) for 2 hours at RT. After blocking, the membrane was cut between the 42 and 21 kDa bands. The primary antibodies (MGMT mouse antibody and Actin rabbit antibody) were diluted in 10 mL blocking buffer (1:100 and 1:200 respectively). Membrane pieces were incubated overnight at 4 °C with their respective antibodies. The next day, the membranes were washed 3 times for 7 minutes with TBS-T. The secondary antibodies (goat anti-rabbit and goat anti-mouse) were also diluted in 10 mL blocking buffer (1:200 and 1:2000 respectively) and added on the membranes for 2 hours at room temperature protected from light. Afterwards, membranes were again washed 3 times for 7 minutes with TBS-T. For the secondary antibody Goat anti-mouse horse radish peroxidase antibody, ECL Western substrate reagents were mixed 1:1 and added to the membrane. Goat anti-rabbit Alexa Fluor 488 antibody was added to the membrane. Fluorescent and chemiluminescent images were taken using a ChemiDoc™ MP Imaging System.

#### Cell exposure to TMZ for DNA extraction

LN-229 cells (WT and +MGMT) were seeded in 10 cm dishes (1 x 10^6^ cells per dish) and incubated for 24h. Cells were exposed to increasing TMZ concentration (50 µM to 1 mM TMZ or 3x 1 mM TMZ, 1% DMSO for all conditions) or solvent control (1% DMSO) in normal growth medium. After 24 hours, cells were detached with Trypsin and genomic DNA was extracted using the QIAamp DNA Mini kit according to the manufacturer’s protocol.

#### Naked genomic DNA exposure to TMZ

Purified genomic DNA from LN-229 cells in Tris-HCl buffer (10 mM, pH 7.4) was incubated with 1 mM TMZ (1% DMSO final) on a thermoshaker at 37 °C and 1100 rpm for 24h. The reaction mixture was purified by ethanol precipitation.

#### *O*^6^-MeG quantification

*Reagents.* Phosphodiesterase I from *Crotalus adamanteus* venom (Sigma, P3243), Benzonase nuclease (Sigma, E1014) and Alkaline phosphatase from bovine intestinal mucosa (Sigma, P5521) were used for sample digestion. Samples were purified with Polyethersulfone (PES) 10kDa MWCO filters (VWR, 516-0230P) and Sep-Pack Vac C18 1cc/50mgv columns (Waters, WAT054955) or Strata™-X 33 mm Polymeric Reversed Phase columns (30mg/mL, Phenomenex, 8B-S100-TAK). *O*^6^-methyl-d_3_-deoxyguanosine (*O*^6^-Me-d_3_-dG) was purchased from Toronto Research Chemicals. *O*^6^-methyl-deoxiguanosine (*O*^6^-me-dG), ammonium hydroxide (ACS Reagent 28–30%), hydrochloric acid (ACS Reagent 37%), and acetic acid (HPLC grade) were purchased from Sigma. To concentrate or dry samples, a miVac Duo Centrifugal Concentrator (Genevac™) was used. HPLC analysis was carried out on an Agilent 1100 or 1200 series HPLC system equipped with a Phenomenex Kinetex 2.6 µm C18 100 Å column (2.1×150 mm). Mass spectrometric analysis was carried out on a Waters nanoAcquity UPLC system, with a Phenomenex Luna Omega 3 µm Polar C18 100 Å, 150 x 0.5 mm column, coupled to a Thermo Scientific TSQ Vantage triple quadrupole mass spectrometer.

#### Sample preparation

DNA was hydrolyzed to yield deoxyribonucleosides as previously described.(23) Briefly, phosphodiesterase I from *Crotalus adamanteus* venom (0.03 U/10 µg DNA), benzonase nuclease (25 U/10 µg DNA) and alkaline phosphatase from bovine intestinal mucosa (20 U/10 µg DNA) were mixed in 50 µL digestion buffer (20 mM Tris-HCl, 100 mM NaCl, 20 mM MgCl_2_, pH 7.6) and added to dried DNA. Each sample was spiked with 0.2 pmol of deuterium labelled standard *O*^6^-me-d3-G. Samples were incubated at 37 °C for 6 h. After the incubation, 450 µL of H_2_O was added to reach 500 µL, and digestion enzymes were removed by filtration over a PES 10kDa MWCO filter. An aliquot of the resulting solution (50 µL) was reserved for quantification of 2’-deoxyguanosine (dG). For the remaining solution, modified nucleosides were enriched by solid-phase extraction using Sep-Pack Vac C18 1cc/50mgv columns or Strata™-X 33 mm Polymeric Reversed Phase columns. First, the columns were washed twice with 1 mL methanol and equilibrated twice with 1 mL H_2_O, then the samples were added. Wash steps included twice 1 mL H_2_O and 1 mL 3% methanol, and samples were eluted twice with 450 µL 80 % methanol. Using the Phenomenex columns, there was an additional wash step with 10% methanol and the samples were eluted in 1 mL 50% methanol. Samples were then dried in conical glass inserts. The dried samples were frozen at -20 °C and resuspended in 4 or 10 µL H_2_O prior to measurement.

#### dG analysis on HPLC

The dG content of enzymatically digested DNA samples was determined by HPLC with UV-detection at 254 nm using a 20 μL aliquot of the digestion mix. The nucleosides were separated on an *Agilent 1100 or 1200 series* HPLC system equipped with a *Phenomenex Kinetex 2.6 µm C18 100 Å* column (2.1×150 mm). Two different methods were used for the time course and the TMZ dose range experiments respectively. The first one included applying a gradient starting at 3% acetonitrile in H_2_O [A] followed by increasing proportions of acetonitrile [B] at a flow rate of 200 μL/min: 0→6.0min, 0→15% (v/v) B; 6.0-6.1min, 15→80% (v/v) B; 6.1→11.0min, 80% (v/v) B; 11.0→11.2min, 80→0% (v/v) B. The second method included applying a gradient starting at 100% H_2_O [A] followed by increasing proportions of acetonitrile [B] at a flow rate of 400 μL/min: 0→5.0min, 0→10% (v/v) B; 5.0-8.0min, 10→100% (v/v) B; 8.0→9.0min, 100→0% (v/v) B. Additional 15 minutes were used to re-equilibrate the column in [A] for both methods. Calibration curves were made by analysis of dG in H_2_O in six concentrations in the range of 0.1 μM – 50 μM and injection volumes of 20 μL. The dG concentration in each 20 μL digestion mix aliquot was determined from the calibration curve and used to estimate the total number of nucleotides (#nt) in each sample, assuming a GC content of 41% in the human genome according to the following equation (1) where V_tot_ (L) is the total volume of the digestion mix and N_A_ is the Avogadro constant (6.022*10^23^ mol^-1^).

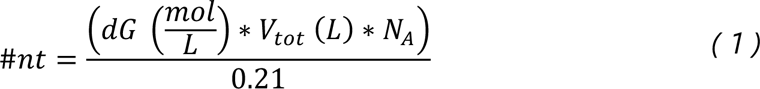

#### Mass spectrometric measurement of O^6^-me-dG

Stable isotope dilution LC-ESI-MS/MS experiments for detection and quantification of *O*^6^-me-dG were performed using a *Waters nanoAcquity UPLC* system coupled to a *Thermo Scientific TSQ Vantage* triple quadrupole mass spectrometer. *A Phenomenex Luna Omega 3 µm Polar C18 100 Å, 150 x 0.5 mm* column, maintained at 40 °C, was used for analysis. The samples were kept at 12°C in the autosampler until analysis. A gradient starting at 100% of 0.1% formic acid in H_2_O [A] followed by increasing proportions of 0.1% formic acid in acetonitrile [B] up to 90%, at a flow rate of 10 µL min^−1^ over 30 minutes. An additional 20 minutes were used to wash and re-equilibrate the column under the starting conditions. The sample injection volume was set to 2 μL. The mass spectrometer was operated in positive electrospray ionization mode and *O*^6^-me-dG was analyzed in selected reaction monitoring mode (SRM) as its [M+H]^+^ species. The general source-dependent parameters were as follows: capillary temperature 250 °C, spray voltage 3000 V, sheath gas pressure 30 (arbitrary unit), and collision gas pressure 1 mbar. Tuned S-lens values were used, and the scan width and scan time were set to 0.1 m/z and 0.1 s, respectively. The parent mass (m/z) is 282.3 and 285.3 for *O*^6^-me-dG and *O*^6^-me-d3-dG respectively, while the fragment mass is 166.1 and 169.1. Collision energy is 16 V for both. Calibration lines were prepared in H_2_O in the range of 0.25 – 50 nM or 4 – 200 nM *O*^6^-me-dG using 7 calibration points spiked with a final concentration 25 or 20 nM *O*^6^-me-d3-dG internal standard each. Data was processed using Thermo Xcalibur Quanbrowser (Ver. 2.1.0.1139) using the internal calibration method. The total number of *O*^6^-me-dG lesions (#O^6^ medG) in each sample was determined from the calculated concentration according to the equation (2):

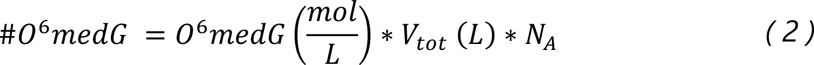

where *O*^6^medG (mol/L) = *O*^6^-me-dG concentration determined via LC-MS/MS-detection, V_tot_ (L) = total volume of mass spec sample, N_A_ = Avogadro number = 6.022*10^23^ mol^-1^.

### *O*^6^-MeG-seq

#### Reagents

High fidelity polymerases Vent® (exo-) DNA Polymerase (M0257S), Deep Vent® DNA Polymerase (M0258S) and Q5® High-Fidelity DNA Polymerase (M0491S) as well as other reagents for library preparation NEBNext® Ultra™ II DNA Library Prep Kit for Illumina® (E7645S), NEBNext® Ultra™ II Q5® Master Mix (M0544S), Instant Stick-end Ligase Master Mix (M0370S), Exonuclease I (M0293S), Q5® Reaction Buffer (B9027S), ThermoPol® Reaction Buffer (B9004S), Deoxynucleotide (dNTP) Solution Sets (N0446S) and NEBNext® Multiplex Oligos for Illumina® (E7335S) were purchased from New England Biolabs. Platinum™ SuperFi II DNA Polymerase (Invitrogen™, 12361250), dNTP Set 100 mM Solutions (R0181) and Pierce™ enhanced chemiluminescence (ECL) Western Blotting Substrate (32109) were purchased from Thermo Scientific™. For immunoprecipitation Rabbit Anti-Mouse IgG H&L (Abcam, ab46540) and anti-*O*^6^-me-dG antibody (Squarix, EM 2-3, SQM003.1) were used in addition to Dynabeads™ Protein G for Immunoprecipitation (Invitrogen™, 10003D), Dynabeads™ M-280 Sheep Anti-Rabbit IgG (Invitrogen™, 11203D), Dynabeads™ MyOne™ Streptavidin C1 (Invitrogen™, 65001) and sheared salmon sperm DNA (Invitrogen™, AM9680), which were purchased from Thermo Scientific™. DNA was purified with AMPure XP DNA purification beads (Beckman Coultier, A63880) and concentration was measured using the Quantus™ Fluorometer (Promega, E6150) with QuantiFluor® ONE dsDNA dye (Promega, E4870). Rotating incubation was done using a Tube Revolver Rotator from Thermo Scientific™ (88881002).

#### Oligonucleotides used in the primer extension assay

All oligonucleotides were synthesized and HPLC-purified by Eurogentec. The primer extension system consisted of a Cy3 labelled 25 mer primer, 5’-Cy3-ATA GGG GTA TGC CTA CTT CCA ACT C-3’ and a 40 mer template 5’-GAG GTG AGT TXA GTG GAG TTG GAA GTA GGC ATA CCC CTA T-3’ (X = G, *O*^6^-meG, 8-oxoG or tetrahydrofuran) or the same sequence with *O*^6^-MeG context GXC and TXT instead. Cy3 labelled oligonucleotides were used as markers for the 40 mer full length, 5’-Cy3-GAG GTG AGT TGA GTG GAG TTG GAA GTA GGC ATA CCC CTA T-3’ and a 29 mer for the stalling site, 5’-Cy3-ATA GGG GTA TGC CTA CTT CCA ACT CCA CT-3’. All oligonucleotides were diluted in H_2_O. The 40 mer templates (1.5 µM final concentration) were annealed with the 25 mer primer (1 µM final concentration) by heating the mixture to 95 °C, followed by a slow cool-down to room temperature over 4 h.

#### Primer extension assay

The capacity of polymerases to bypass *O*^6^-meG was assessed by primer extension assays. Reaction mixtures contained the 40mer template previously annealed with the primer (0.1 µM), dNTPs (200 µM), 1X ThermoPol buffer (when using Vent or Deepvent) or 1X Q5 buffer (when using Q5), and polymerase (Vent, DeepVent or Q5, 0.4 U final) and topped up with H_2_O to a final reaction volume of 20 µL. For SuperFi, the mixture consisted of the 40mer template previously annealed with the primer (0.1 µM) and SuperFi II 2X mastermix topped up with H_2_O to a final reaction volume of 20 µL. The primer extension reaction was performed in a thermocycler for 10 min using the optimal temperatures for each polymerase: 75 °C for Vent, 72 °C for DeepVent, 65 °C for Q5, or 72 °C for SuperFi II. Reactions were stopped by adding 40 µL of a quenching solution (80% formamide, 0.5 M NaOH, 0.5 M EDTA and bromophenol blue). Samples (10 µL) were loaded on a 20% acrylamide containing 7 M urea denaturing gel and run with TBE running buffer (0.9 M Tris base, 20 mM EDTA and 0.9 M boric acid) for 1 h at 120 V. Extension products were imaged using a BioRad imager with Cy3 settings.

#### Oligonucleotides used in library preparation

AD1T: 5’-phos-GAT-CGG-AAG-AGC-ACA-CGT-CTG-AAC-TCC-AGT-CA-SpC3; AD1B: 5’-NNN-NNG-ACT-GGT-TCC-AAT-TGA-AAG-TGC-TCT-TCC-GAT-C*T (* indicating a phosphorothioate bond); AD2T: 5’-phos-AGA-TCG-GAA-GAG-CGT-CGT-GTA-GGG-AAA-GAG-TGT-SpC3; AD2B: 5’-ACA-CTC-TTT-CCC-TAC-ACG-ACG-CTC-TTC-CGA-TCT-NNN-NN-SpC3; O3P: 5’-biotin-GAC-TGG-AGT-TCA-GAC-GTG-TGC-TCT-TCC-GAT-CT; and SH: 5′-biotin-NNG-ACT-GGT-TCC-AAT-TGA-AAG-TGC-TCT-TCC-G-SpC3. AD1 and AD2 adaptors were prepared by mixing equal volumes (20 µL) of 100 µM AD1T/AD2T and AD1B/AD2B with 10 µL 5X annealing buffer (50 mM Tris, pH 8.0, 250 mM NaCl, 5 mM EDTA) and heating to 98 °C, then slowly cooling to 25 °C over 4 h. Oligonucleotides and indexing primers were synthesized and HPLC-purified by Eurogentec.

#### O^6^-MeG-seq library preparation

For library preparation, 1.5 μg of DNA was sheared using Q800 sonicator from Qsonica to produce fragments having an average length of 400 bp using the following settings: 20%, 3 min, 2 s on/ 5 s off. DNA fragments <200 bp in length were removed by size-selective purification with AMPure XP beads (1:1 v/v beads:DNA) resulting in a size-range of about 200-1000 bp. DNA concentration was measured, and 900 ng were used for further library preparation. DNA was end repaired and AD1 (40 µM) was ligated according to the instructions of NEBNext® Ultra™ II DNA Library Prep Kit for Illumina®. The ligation mixture was incubated at 4 °C overnight. The ligation product was purified with AMPure XP beads (0.7:1 v/v beads:DNA) and eluted with 12 μL 0.1X TE buffer. Eluted DNA was denatured by mixing it with urea (5 μL, 8 M stock), heating (98 °C, 2 min), and immediately cooling it on ice. The denatured DNA was mixed with 2.5 µL of 8X IP buffer (160 mM Tris-HCL, 1.2 M NaCl, pH 7.5, 4% Triton X-100, 4 °C), and antibody-coated beads.

The antibody-coated beads were prepared ahead of use by mixing 1.25 μL of protein G Dynabeads and 1.25 μL anti-rabbit Dynabeads. The beads were washed twice using 100 μL 1X IP buffer (20 mM Tris-HCL, 150 mM NaCl, pH 7.5, 0.5% Triton X-100, 4 °C). Then 4.5 μL 1X IP buffer, 0.25 μL salmon sperm DNA, 0.5 μL rabbit anti-mouse IgG, and 0.5 μL anti-*O*^6^-meG antibody was added to the beads. The beads were resuspended by pipetting, and they were incubated at 4 °C overnight rotating with oscillation on a tube revolver to allow for binding of the complementary antibodies. The resulting antibody-coated beads were washed using 1X IP buffer (100 μL, 4 °C), resuspended in 5 mL 1X IP buffer and 0.5 μL of salmon sperm DNA, and mixed with the DNA solution described above. The mixture was resuspended and incubated at 4 °C overnight rotating with oscillation on a tube revolver to allow for binding of the antibodies to *O*^6^-meG in the DNA. The beads, now bound to *O*^6^-meG-containing DNA fragments, were washed three times with 180 μL 1X IP buffer and once with 1X TE buffer, then eluted twice with 50 μL elution buffer (10 mM Tris-Cl pH 8.0, 1 mM EDTA, 1% SDS, 65 °C) at 65 °C, 1100 rpm for 5 min. The combined elution fractions were purified by phenol-chloroform extraction followed by ethanol precipitation and was then resuspended in 6 μL 0.1X TE buffer.

The purified DNA was then mixed with 1.5 μL of O3P primer (20 μM) and 7.5 μL of SuperFi II 2X mastermix. Primer extension was performed under the following conditions: 50 s at 98 °C, 5 min at 72 °C, and hold at 37 °C. Exonuclease I (30 U) was added to the resulting mixture, which was incubated at 37 °C for 15 min to digest the excess primer that was not extended. The resulting mixture was purified with AMPure XP beads (37 µL beads, 25 µL H_2_O, corresponding to 0.9:1 v/v beads:DNA) and eluted with 20 μL 0.1X TE buffer. The purified DNA was then mixed with 2 μL of the biotinylated SH primer (10 μM stock), 25 μL 1 B&W buffer (5 mM Tris-HCl pH 8.0, 0.5 mM EDTA, 1 M NaCl, 0.1% Tween20, 0.1%CA-630), and subjected to a slow annealing process using a thermocycler programmed with the following conditions: 2 min at 98 °C, then cooling with 1 min/°C from 97 °C to 76 °C, 5 min/°C from 75 °C to 55 °C, 1 min/°C from 54 °C to 25 °C, and hold at 4 °C. To capture the SH oligo, 10 μL Dynabeads MyOne Streptavidin C1 were washed twice with 1X B&W buffer, resuspended with 5 μL 5X binding buffer (50 mM Tris-HCl pH 8.0, 5 mM EDTA, 2.5 M NaCl, 0.1% Tween20, 0.1% CA-630, 25 mM MgCl_2_) and added to the annealing product. The mixture was incubated for 1 h at 4 °C rotating with oscillation on a tube revolver. After incubation, the supernatant containing the desired DNA was transferred to a new 1.5 mL tube, and the beads were washed with 50 µL 1X B&W buffer. The supernatants were combined and purified by ethanol precipitation. Sodium acetate was not added due to the high salt concentration in the supernatant. The air-dried pellet was resuspended in 6.5 μL 0.1X TE buffer. The purified DNA was denatured for the following AD2 ligation by heating to 98 °C for 2 min and then immediately cooled on ice. For the ligation, 1 μL AD2 (40 μM stock) and 7.5 µL of Instant Sticky-end Ligase Master Mix (2X stock) were added and incubated overnight at 4 °C. The DNA was purified with AMPure XP beads (40 µL beads, 35 µL H_2_O, corresponding to 0.8:1 v/v beads:DNA) and eluted with 16 μL 0.1X TE buffer. 0.3 mL were amplified with specific and unspecific primers and the products were run on a 5% neutral-PAGE. The rest was amplified using NEBNext Ultra II Q5 Master Mix with dual indexing primers for Illumina. The amplified products were purified by AMPure XP beads (0.9:1 v/v beads:DNA), and DNA concentration was determined. The libraries were pooled and purified again by AMPure XP beads (0.9:1 v/v beads:DNA) to remove residual primer-dimers, then eluted using 10 mM tris buffer (pH 8.0). The pool was diluted to 10 nM for sequencing. The pooled libraries were sequenced on a NovaSeq Illumina sequencer (single end) by the Functional Genomic Center of Zürich.

#### Data availability

*O*^6^-meG-seq read files can be found on NCBI Gene Expression Omnibus (GEO) (accession number GSE249155). Other data and support files can be found on Zenodo (DOI 10.5281/zenodo.10518966). Data analysis scripts and notebooks are also available at gitlab.ethz.ch/eth_toxlab/o6meg-seq.

#### Sequencing data processing and analysis

After demultiplexing of sequencing data, each sample was represented by a fastq.gz file containing 101-nucleotide-long genomic reads. The quality of the raw sequencing data was checked via FastQC version 0.11.9. Low-quality reads and adapter-containing reads were removed via trimmomatic version 0.38 with the following parameters: SE ILLUMINACLIP:Trimmomatic-0.39/adapters/TruSeq3-SE.fa:2:30:10 LEADING:3 TRAILING:3 SLIDINGWINDOW:4:15 MINLEN:101. Reads containing the AD1B sequence, which were not removed by subtractive hybridization, were discarded using cutadapt and the following settings: GACTGGTTCCAATTGAAAGTGCTCTTCCGATCT; e=0.1; min_overlap=15. The reads were mapped to human reference genome GRCh38 via bowtie2 version 2.3.5.1, using pre-built bowtie2 index from https://genome-idx.s3.amazonaws.com/bt/GRCh38_noalt_as.zip and applying otherwise standard settings. Read duplicates were removed by gatk (version 4.2.0.0) MarkDuplicatesSpark. Samtools version 1.12 were employed to sort, index, and generate statistics of bam files. Bedtools2 version 2.29.2 were used to retrieve the coordinates of mapped and deduplicated reads and extract the sequence context of the modified nucleotides from the reference genome, while the -1 position of the read start was the modification site. The downstream analysis of DNA-modification data and their visualization were performed via custom scripts in Python with indicated modules in Jupyter notebooks, which can be found on the data repository (DOI 10.5281/zenodo.10518966) and at gitlab.ethz.ch/eth_toxlab/o6meg-seq. For signature extraction, data was processed using the R package MutationalPatterns (https://github.com/UMCUGenetics/MutationalPatterns). For the analysis of whole-genome distributions and in relation to gene expression and chromatin accessibility, only data from GRCh38’s chromosome 1-22 and chromosome X was used.

#### Data Resources

The following public datasets were employed in the data processing and analysis: GRCh38 was downloaded from NCBI (https://ncbi.nlm.nih.gov/projects/genome/guide/human/index.shtml). COSMIC single base substitution signatures from COSMIC_v3.3.1_SBS_GRCh38.txt (https://cog.sanger.ac.uk). Centromere and gap coordinates were obtained from UCSC Table Browser (https://genome.ucsc.edu/). ATAC-seq data was downloaded from Chip-atlas.org (https://chip-atlas.org/peak_browser searching for ATAC-Seq and LN-229). Transcript coordinates, GENCODE/V41/knownGene, Canonical transcripts of genes, GENCODE/V41/knownCanonical, and Protein-coding genes, GENCODE/V41/knownToNextProt, were obtained from UCSC Table Browser. Genes were represented by canonical transcripts between transcription start site (TSS) and transcription end site (TES). Gene expression data, OmicsExpressionProteinCodingGenesTPMLogp1, was obtained from DepMap Public 23Q2 (https://depmap.org/portal/download/all/), the cell-line accession number ACH-000595 (LN-229). Oncogenes were obtained from COSMIC Cancer Gene Census, Tier 1 (https://cancer.sanger.ac.uk/census, downloaded on 15.05.2023).

#### Statistical analysis

Spearman correlation was used to compare replicates with each other and compare our data to existing ATAC-seq data. Cosine similarity was used to compare extracted *O*^6^-meG signatures with COSMIC SBS signatures. Wilcoxon test was performed using Python’s module scipy version 1.6.3. All calculations can be reviewed in the Jupyter notebooks at gitlab.ethz.ch/eth_toxlab/o6meg-seq.

## RESULTS

### *O*^6^-MeG stalls SuperFi II polymerase as a basis for marking its location in DNA

To map *O*^6^-MeG in genomic DNA, we established a new method termed *O*^6^-MeG-seq (Figure 1A). Following antibody-induced capture of *O*^6^-MeG-containing DNA fragments, specific marking of adduct locations in related modification-mapping methods (17,18,20) requires that the adduct induces stalling of a DNA polymerase. However, O^6^-MeG is easy for most polymerases to bypass. To identify a suitable polymerase enzyme for this mapping application, we tested the capacity of four different high-fidelity polymerases to bypass *O*^6^-MeG present at position 29 in a 40mer oligonucleotide. Amongst Vent, DeepVent, Q5 and SuperFi II, only SuperFi II polymerase was stalled at position 29, resulting in a truncated product and a negligible indication of full-length product (Supp. Figure 1). We tested the process in additional trinucleotide contexts flanking *O*^6^-MeG in the template and found the same stalling effect, suggesting it is likely to occur regardless of local sequence differences. Finally, we also tested whether SuperFi-II is stalled by other common DNA modifications such as 8oxoG or abasic (AP) sites, which can result from depurinated-TMZ-induced N7-methylguanine that are about ten times more abundant after TMZ exposure than *O*^6^-MeG.(6) We found that 8oxoG partially stalled and tetrahydrofuran (THF, a stable analog of an AP site) completely stalled the polymerase (Supp. Figure 2). This means that although fragments containing-*O*^6^-MeG are enriched using an antibody, AP sites or 8-oxoG, could nevertheless still be present and stall SuperFi II, as a basis of possible artifacts that are called as *O*^6^-MeG sites in the sequencing data. Indeed, a common limitation to the use of polymerase stalling as a strategy for marking DNA modifications is the difficulty to identify clustered modifications since only the first stall per strand is marked.

**Figure 1.**
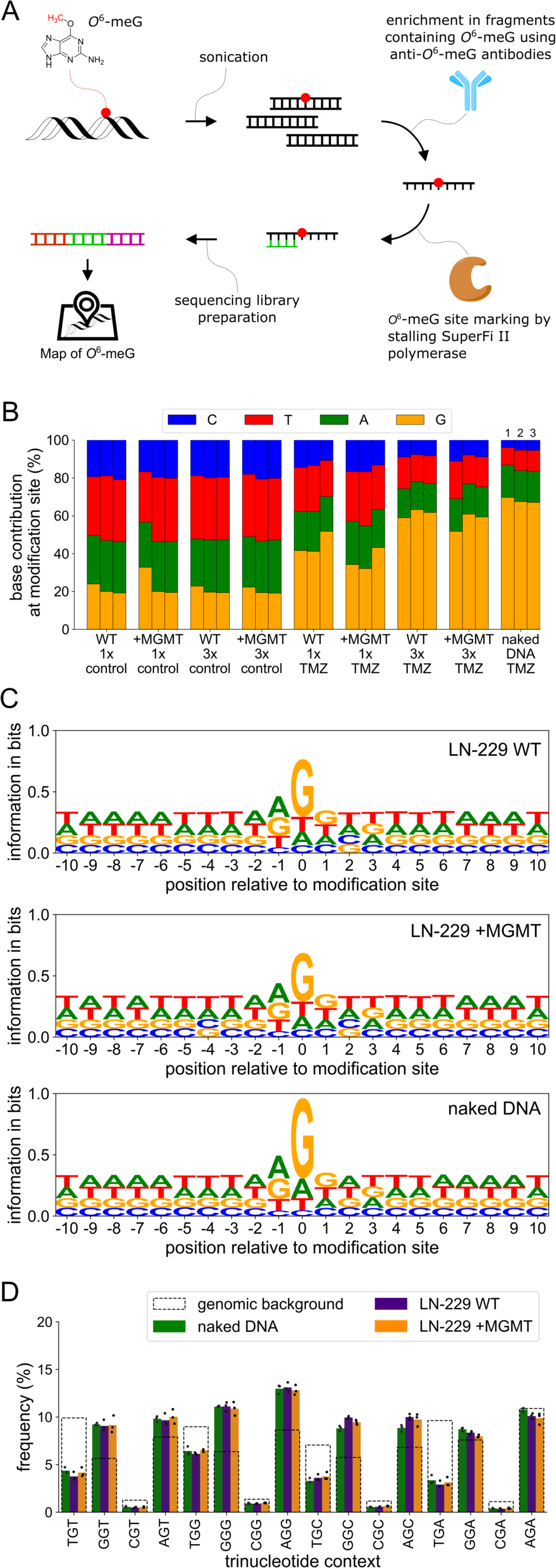
*O^6^-MeG is induced in distinct trinucleotide patterns by TMZ*. **A** Strategy for *O*^6^-MeG-seq. DNA fragments containing *O*^6^-MeG are pulled down with an *O*^6^-MeG specific antibody. The exact site is marked with SuperFi II polymerase stalling at *O*^6^-MeG. Illumina library preparation and sequencing are used to map *O*^6^-MeG at single-nucleotide resolution. **B** Base contribution at modification site. 3 biological replicates of solvent control or TMZ-exposed LN-229 cells (WT or +MGMT) and TMZ-exposed naked DNA from different genomic DNA sources: 1: LN-229 WT (not transfected), 2: LN-229 WT (transfection control), 3: LN-229 +MGMT. **C** Position information across 10 bases upstream and downstream of modification sites in 3x 1 mM TMZ-exposed LN-229 cells and TMZ-exposed naked DNA. The relative heights of the letters corresponding to bases indicate their relative abundance at that site, while the height of the stack is defined by the certainty these bases are found at this position (max. 2 bits for 4 bases). **D** Trinucleotide context frequencies of *O*^6^-MeG considering only guanines at modification site.

### MGMT partially repairs TMZ-induced *O*^6^-MeG in LN-229 +MGMT cells

As a relevant cell model to study the influence of MGMT on *O*^6^-MeG distribution throughout the genome, we selected LN-229 cells, which originate from a female glioblastoma patient, have low MGMT activity, and are responsive to TMZ.(24) Thus, to probe the impacts of MGMT expression on *O*^6^-MeG levels and locations, we used LN-229 cells transfected with an *MGMT* harboring plasmid (+MGMT) and a transfection control, referred to as LN-229 wild type (WT) for clarity. We confirmed by western blot that LN-229 WT cells did not express MGMT, and the LN-229 +MGMT cells highly expressed MGMT (Supp. Figure 3). To benchmark the sensitivity of the cells to TMZ and determine corresponding *O*^6^-MeG levels, cells were exposed to increasing concentrations of TMZ (50 µM – 1 mM) and repetitive exposure (3x 1 mM TMZ) for up to 6 d and their viability and *O*^6^-MeG levels were assessed at different time points between 0 and 144 h. Viability in LN-229 WT cells decreased more compared to LN-229 +MGMT cells three and six days after exposure to TMZ (Supp. Figure 4). For both cell lines, viability was not affected 24 h after exposure except for repeated exposure to 1 mM TMZ, where the viability was slightly reduced in both cell lines. Peak *O*^6^-MeG levels were observed 24 h after exposure to TMZ (Supp. Figure 5A). In all conditions, there is roughly two-fold more *O*^6^-MeG in LN-229 WT vs. LN-229 +MGMT cells (Supp. Figure 5B).

### *O*^6^-MeG is induced in distinct trinucleotide patterns upon TMZ exposure

To test the hypothesis that *O*^6^-MeG trinucleotide patterns are similar to SBS 11 found in patients previously treated with TMZ (25), we characterized the relative frequency of *O*^6^-MeG occurrence in different trinucleotide contexts and compared these to all COSMIC mutational signatures.(26) A benefit of *O*^6^-MeG-seq is that it is possible to map *O*^6^-MeG locations at single-nucleotide resolution, thereby determining their relative frequencies in a sequence context manner. Since the polymerase stalls before *O*^6^-MeG, the -1 position of the read start site is the modification site. We exposed LN-229 WT and LN-229 +MGMT cells to 1 mM TMZ once and three times within a 24 h period and prepared sequencing libraries for *O*^6^-MeG mapping. We prepared three biological replicates for every exposure condition in addition to the corresponding solvent controls. As positive control, naked DNA extracted from LN-229 cells was exposed to 1 mM TMZ for 24 h, which resulted in high amounts of *O*^6^-MeG (3,300 to 8,200 *O*^6^-MeG per 10^7^ nt). Unfortunately, it was not possible to prepare sequencing libraries from cells exposed to clinically relevant concentrations of up to 100 µM TMZ (27,28), as there was insufficient *O*^6^-MeG in the DNA (below 100 *O*^6^-MeG per 10^7^ nt, Supp. Figure 5), resulting in inadequate DNA yield during immunoprecipitation for library preparation. Guanine was enriched at the modification site in the genomes from cells and naked DNA exposed to 1 mM TMZ, while solvent control samples reflect nucleotide proportions of the human genome (29) (Figure 1B).

To analyze base proportions in the vicinity of the modification site, information content was calculated from a total of 21 nucleotides at and around the modification site and plotted as logos with information in bits.(30) The relative heights of the letters corresponding to bases indicate their relative abundance at that site, while the height of the entire stack of letters reflects deviation from randomness at this position with a maximum of 2 bits. We found that nucleotide ratios were altered only for bases directly flanking *O*^6^-MeG (Figure 1C). Most significantly, *O*^6^-MeG formed preferentially 3’ of guanine and adenine since their frequency was higher than expected from the genomic background (Figure 1D). Additionally, there were slightly more *O*^6^-MeG sites 5’ of cytosine in both cell lines compared to exposed naked DNA. There were no notable differences in trinucleotide patterns in the LN-229 +MGMT cells compared to LN-229 WT.

### *O*^6^-MeG as precursor of TMZ-related mutational signatures

Analogous to the extraction of mutational signatures (25), we used non-negative matrix factorization of the *O*^6^-MeG-seq trinucleotide patterns to extract *O*^6^-MeG signatures arising from TMZ exposure as putative precursors of mutational signatures. While mutational signatures have 96 features corresponding to the trinucleotide context for each of the 6 possible single base substitutions, DNA modification signatures have only 64 features describing all trinucleotide context possibilities of any base modification.(17) We found two distinct DNA modification signatures designated as A and B (Figure 2A). While signature B resembled the genomic background, in signature A, there were higher *O*^6^-MeG frequencies in the XGY contexts, as expected due to guanine enrichment in TMZ-exposed samples. *O*^6^-MeG-seq trinucleotide patterns from TMZ-exposed cells mainly contributed to signature A while trinucleotide patterns from unexposed controls contributed to signature B (Figure 2B). The *O*^6^-MeG signatures were then compared to all COSMIC SBS signatures using the cosine similarity metric (Figure 2C). As the signatures cannot be directly compared due to their different dimensions, the XGY contexts of the *O*^6^-MeG signatures were converted into X’[C>T]Y’, X’[C>A]Y’, X’[C>G]Y’, X’[T>A]Y’, X’[T>C]Y’ or X’[T>G]Y’ mutations, where X’ and Y’ are reverse complements of the flanking bases present in the modified triplets. All other contexts were set to zero. As an example, XGY were converted to X’[C>T]Y’, while all other base substitutions, i.e. X’[C>A]Y’, X’[C>G]Y’, X’[T>A]Y’, X’[T>C]Y’ and X’[T>G]Y’, were zero. This converted signature was then compared to COSMIC SBS signatures. Only when XGY from signature A was converted to X’[C>T]Y’ was there a high similarity (cosine similarity ≥ 0.9) to any mutational signatures. This made sense since *O*^6^-MeG is known to cause mostly C to T mutations.(11) Moreover, the similar signatures were SBS 11, which has been found in cancer tissue of patients previously treated with TMZ (25), and SBS 23, which has so far not been linked to an aetiology (Figure 2C,D and Supp. Figure 6).(26,31)

**Figure 2.**
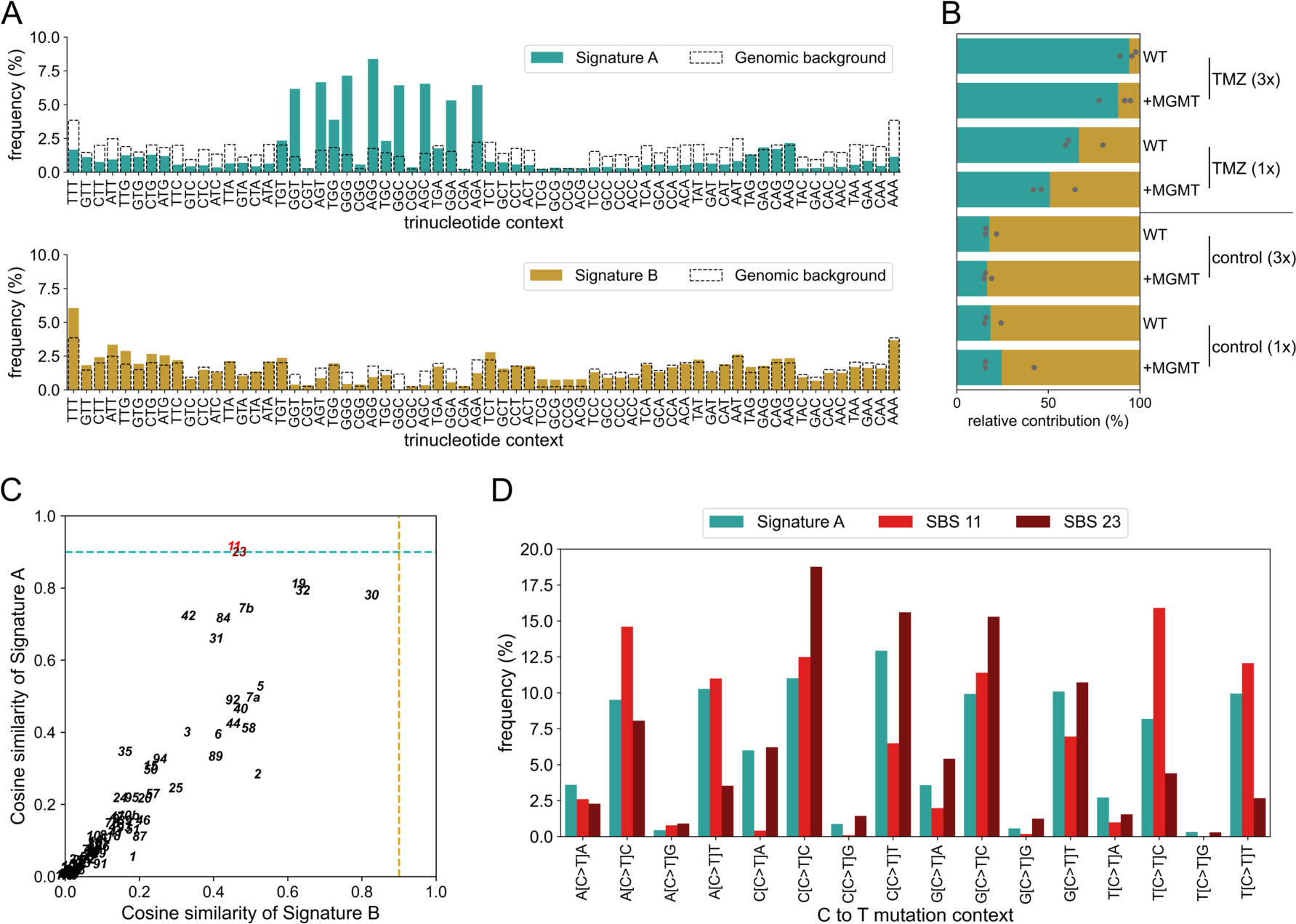
*O*^6^-MeG as precursor of TMZ-related mutational signatures. **A** Extracted *O*^6^-MeG signatures, termed signature A and B. Signatures were extracted using non-negative matrix factorization. **B** Relative contribution of samples to the extracted signatures A and B. Three biological replicates per sample condition. Control samples were exposed to 1% DMSO. **C** Cosine similarities of all COSMIC SBS compared to both *O*^6^-MeG signatures. Conversion of *O*^6^-MeG trinucleotide contexts considers reverse complementary trinucleotide contexts of guanine converted into C to T mutations and assumes no signal for other single base substitutions. Cosine similarity of 0.9 was used as cut off for high similarity (dashed lines). **D** C to T mutation contexts of *O*^6^-MeG signature A and COSMIC SBS 11 and 23.

### MGMT does not influence *O*^6^-MeG distribution in the human genome

Having established a characteristic *O*^6^-MeG signature for TMZ, we were interested to compare the genome-wide distribution of *O*^6^-MeG to genome annotations, in order to understand how underlying genomic features may impact formation of *O*^6^-MeG and its repair by MGMT. Thus, *O*^6^-MeG-seq data from TMZ-exposed samples were filtered for reads with only guanine at the modification site and binned in 100 Kb bins. The bins were then normalized by guanine-only read depth and bin size (Figure 3A,B) or by guanine-only read depth and genomic guanine abundance per bin (Figure 3C,D). These data were compared with sequencing data analyzed in the same manner, originating from TMZ-exposed naked DNA (Figure 3E,F). In bin-size normalized and G-abundance normalized data, hotspots of *O*^6^-MeG accumulation were found in the genome (Figure 3A-D), however, these seem to be cancelled out when the data from TMZ-exposed cells was compared to the data for TMZ-exposed naked DNA (Figure 3E,F). Additionally, at the resolution of 100 Kb, there appeared to be no difference between LN-229 +MGMT cells and LN-229 WT (Supp. Figure 7A). These findings contradicted our expectations that preferential repair by MGMT might give rise to distinct patterns of *O*^6^-MeG in the genome. Furthermore, we compared whole-genome distributions of *O*^6^-MeG with ATAC-seq data from LN-229 cells (ATAC-seq data was obtained from Chip-atlas.org). *O*^6^-MeG distribution of all samples correlated with ATAC-seq data, with the strongest correlation in the case of higher TMZ exposure (correlation coefficient of 0.3 to 0.5) (Supp. Figure 7B). However, when we compared naked-DNA normalized data with ATAC-seq data, since the influence from chromatin accessibility is only valid in a cell context, there was no correlation.

**Figure 3.**
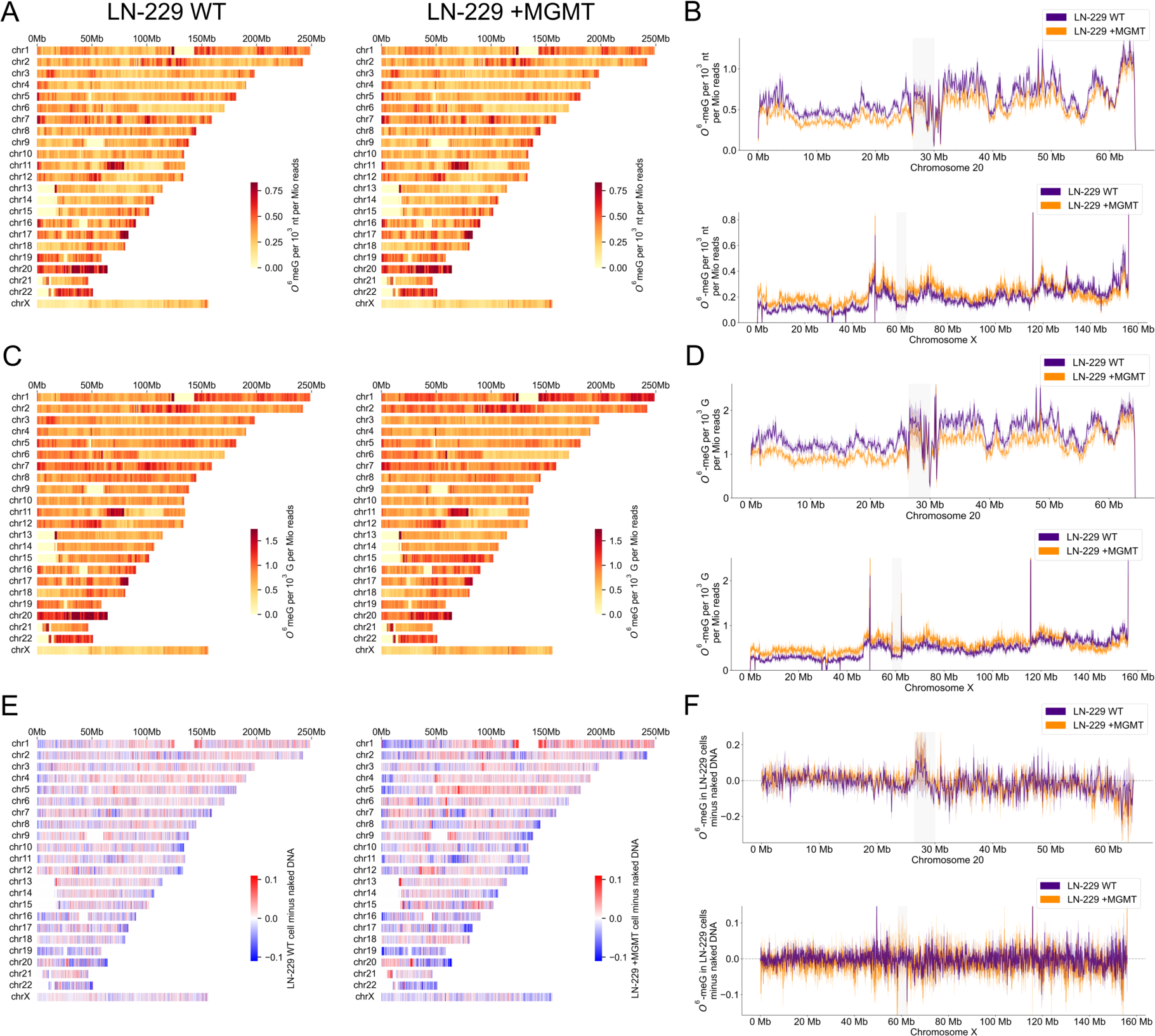
MGMT does not appear to influence *O*^6^-MeG distribution in the human genome. Genome-wide distribution of *O*^6^-MeG in TMZ-exposed LN-229 WT and +MGMT cells. Spearman correlation of replicates was high (>0.8) for all samples used in this analysis (Supp. Figure 8). **A,C,E** Whole-genome view shows the average of 3 biological replicates per bin. Ranges were capped at the 99^th^ and 1^st^ percentile. **B,D,F** *O*^6^-MeG distribution in chromosomes 20 and X of LN-229 WT and +MGMT cells. Faded bands show standard deviation of replicates and centromeric areas were marked by grey background. Ranges were capped at the 99.9^th^ and 0.1^st^ percentile. *O*^6^-MeG abundance was normalized by G-only read depth and bin size (A,B) or by G-only read depth and genomic G abundance (C,D). Additionally, *O*^6^-MeG abundance in cells was corrected by *O*^6^-MeG abundance of TMZ-exposed naked DNA (E,F).

### *O*^6^-MeG has an MGMT-independent strand bias towards the non-transcribed strand in expressed genes while MGMT seems to protect oncogenes

Accumulation of DNA alkylation has been observed to be influenced by gene expression (17), therefore, we analyzed *O*^6^-MeG formation and repair in the context of transcription by comparing the amount of *O*^6^-MeG in transcribed vs. non-transcribed strands of protein-coding genes as a function of their expression (gene expression data was obtained from DepMap Public 23Q2). *O*^6^-MeG counts were normalized by gene length (Figure 4A,B) as well as by G abundance per gene (Figure 4C,D), while in both cases the counts were scaled to the transcribed strand of the unexpressed genes. For both normalizations (gene length and gene G abundance), we observed more *O*^6^-MeG in the non-transcribed strand of highly expressed genes. These patterns were almost identical between the LN-229 WT and +MGMT cells (Figure 4A-D). A distinct non-monotonic behavior was observed as genes in the ≤30% expression tier had more *O*^6^-MeG than genes in the ≤40% tier (Figure 4A-D, right panels). Additionally, we subtracted the *O*^6^-MeG counts in TMZ-exposed naked DNA from the respective *O*^6^-MeG counts in cells (Figure 4E-F), to correct for factors unrelated to gene expression. This correction removed the apparent non-monotonic profile for G-abundance-normalized data. More interestingly, there remained an increasing strand bias for *O*^6^-MeG accumulation with increasing gene expression (Figure 4E,F). This strand bias is apparent only within gene bodies and not in the adjacent upstream and downstream regions (Figure 4G-H). Since this effect is observed in both cell lines, preferential removal in the transcribed strand by MGMT does not seem to be the origin of this phenomenon.

**Figure 4.**
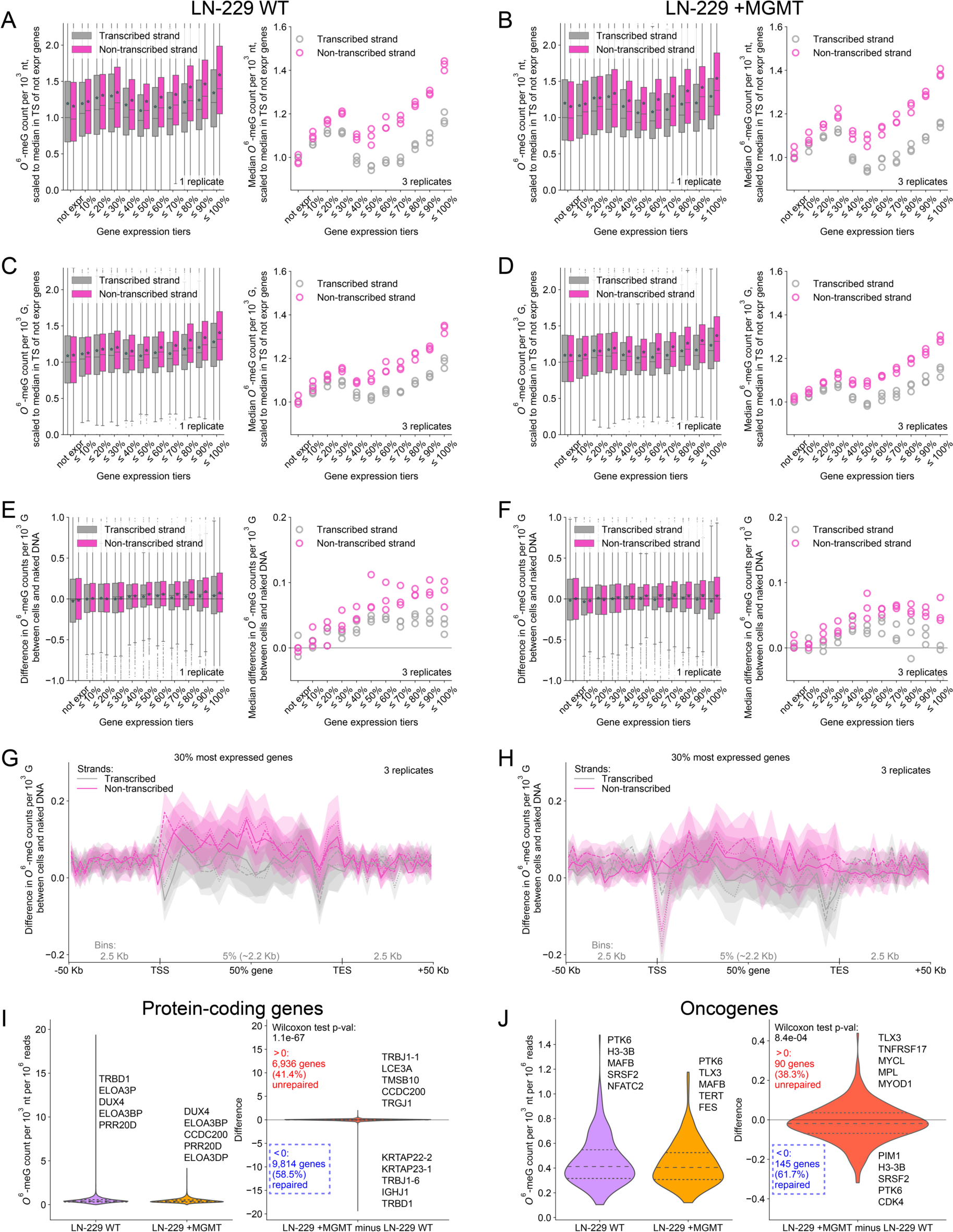
*O*^6^-MeG has an MGMT-independent strand bias towards the non-transcribed strand in expressed genes, but MGMT protects oncogenes from *O*^6^-MeG. **A-F** Gene-specific analysis of *O*^6^-MeG distribution in LN-229 WT (A,C,E) and LN-229 +MGMT (B,D,F) cells with respect to gene expression. Left panels show *O*^6^-MeG abundance in gene expression tiers in one replicate. The asterisks indicate the average abundance. Right panels show the medians of *O*^6^-MeG abundance in gene expression tiers in three replicates together. *O*^6^-MeG abundance was normalized by gene length (A,B) or by gene guanine abundance (C,D). Additionally, *O*^6^-MeG abundance in cells was corrected by *O*^6^-MeG abundance of TMZ-exposed naked DNA (E,F). **G,H** The gene-body profiles of cellular *O*^6^-MeG abundance corrected by *O*^6^-MeG abundance in TMZ-exposed naked DNA. Means and 95% confidence interval of three replicates are shown. TSS: transcription start site. TES: transcription end site. **I,J** *O*^6^-MeG abundance per gene was normalized by read depth, while transcribed and non-transcribed strands were added, and replicates were averaged. Left panels show LN-229 WT and +MGMT while right panels show the difference of LN-229 +MGMT and WT. Analysis was done for all genes (I) or oncogenes only (J).

Because of the mutagenic potential of *O*^6^-MeG, we analyzed *O*^6^-MeG formation and repair in genes regardless of gene expression level and with focus on oncogenes, as their mutations could lead to an increased risk for secondary cancers. For this, *O*^6^-MeG counts were normalized by read depth and gene length. We found the overall *O*^6^-MeG abundance in genes to be lower in LN-229 +MGMT than in LN-229 WT (58.5% of bins have less modifications than in WT) indicating the repair of genes by MGMT (Figure 4I). Furthermore, this difference in *O*^6^-MeG abundance slightly increased when evaluating only oncogenes (61.7% of bins have less modifications than in WT), potentially related to their preferential repair by MGMT (Figure 4J). While these observations were made in a cancer cell line, it suggests that MGMT in normal or stem cells that are also exposed to TMZ during therapy may be better protected, potentially decreasing the risk of secondary cancers caused by TMZ.

## DISCUSSION

By establishing a new method for mapping *O*^6^-MeG at single-nucleotide resolution (*O*^6^-MeG-seq) and combining it with quantitative analysis of alkylation levels, we could elucidate where and to what extent *O*^6^-MeG accumulates in the genome of a glioblastoma cell line upon exposure to the chemotherapeutic drug temozolomide. A key methodological aspect enabling this outcome resided in the discovery that SuperFi II polymerase stalls at *O*^6^-MeG and therefore can be used to mark its location in the genome at single-nucleotide resolution. Therefore, it was possible to analyze trinucleotide contexts of *O*^6^-MeG and identify that certain trinucleotides, particularly GGN and AGN, were preferentially modified by TMZ. Additionally, we could confirm a prominent *O*^6^-MeG signature as a precursor of TMZ mutational signatures. However, *O*^6^-MeG accumulation did not seem to be influenced by particular genomic features, and while MGMT reduces overall levels of *O*^6^-MeG, it does not appear to impact its distribution in terms of sequence contexts or accumulation in genomic features. On the other hand, focusing on *O*^6^-MeG accumulation in genes, we found an MGMT-independent strand bias towards the non-transcribed strand in expressed genes, and less *O*^6^-MeG in oncogenes in LN-229 +MGMT cells than in the WT even after normalizing by read depth to account for overall less *O*^6^-MeG in LN-229 +MGMT.

Levels of *O*^6^-MeG in TMZ-exposed LN-229 WT cells increased in a dose dependent manner from 7 to 1,075 *O*^6^-MeG per 10^7^ nt at drug concentrations ranging from 50 µM to 1 mM TMZ, and there was roughly two-fold more *O*^6^-MeG in LN-229 WT cells than in LN-229 +MGMT cells. The dose-dependent increase of *O*^6^-MeG levels and MGMT-induced decrease is consistent with various earlier studies in different cell lines (27,32–34), confirming that MGMT is effective but incomplete within 24 h in repairing *O*^6^-MeG. Clinically relevant TMZ concentrations up to 100 µM TMZ (33,35) yielded less than 100 *O*^6^-MeG per 10^7^ nt, which were insufficient for mapping by *O*^6^-MeG-seq, therefore, we prepared maps from cells exposed three times to 1 mM TMZ, corresponding to roughly 500 to 1,100 *O*^6^-MeG per 10^7^ nt. While these levels exceed those anticipated to arise from the clinical use of TMZ, it allowed us to reproducibly locate 3.4 – 6.5 Mio *O*^6^-MeG throughout the genome and gain a first genome-wide view on the distribution of *O*^6^-MeG. Additionally, in a previous-study using a similar method for mapping a different type of alkylation damage, namely from benzo(a)pyrene, we did not find a dose-dependent adduct distribution.(17) While this suggests that even maps derived from high concentrations may provide relevant insight, extensive further research is needed to understand how potential distribution changes relative to chemical exposure concentrations, as well as adduct structures and removal mechanisms.

MGMT has been reported to have trinucleotide specificity (36), therefore, it was surprising that there was almost no difference in trinucleotide patterns of *O*^6^-MeG in LN-229 WT and +MGMT cells. Trinucleotide patterns were also similar when naked DNA was allowed to react with TMZ but all patterns in TMZ-exposed samples, cells and naked DNA, were different from the trinucleotide ratios naturally found in the genome (Figure 1D). Since we observed that SuperFi II stalls in different trinucleotide contexts of *O*^6^-MeG, it is unlikely that the observed preference is an artifact of stalling. We interpret therefore, that the favored trinucleotide contexts of *O*^6^-MeG are mostly influenced by adduct formation rather than repair. Further supporting this, as reviewed in Richardson et al. (37), *O*^6^-MeG is preferentially formed 3’ to another guanine, which is in line with our findings that *O*^6^-MeG is preferentially formed 3’ of guanine and adenine (Figure 1D).

Despite the lack of impact of MGMT on the *O*^6^-MeG signature, it was highly compelling that we could link one of the extracted *O*^6^-MeG signatures (signature A) to TMZ exposure by signature contribution analysis of the samples (Figure 2B). Signature A was highly similar to COSMIC SBS 11 (cosine similarity 0.91), found in secondary cancers of patients previously treated with TMZ and was also linked to MMR deficiency(12,38). In comparison, LN-229 cells are MMR proficient (39) and the extracted *O*^6^-MeG signatures from modification maps of TMZ-exposed cells were highly similar to SBS 11. Accordingly, we conclude that SBS 11 can be attributed mainly to TMZ exposure, or other methylating agent with a similar basis of alkylation (40,41). On the other hand, MMR deficiency might be necessary for the accumulation of mutations that manifest as mutational signatures initiated by the chemistry of the methylating agent. This raises the interesting question of what *O*^6^-MeG signature would persist in MMR-deficient cells, which could be further addressed in future work enabled by the methodology established herein.

Analyzing *O*^6^-MeG in the whole genome in 100 Kb bins, we found heterogenous *O*^6^-MeG distributions, however, these did not correspond to known genomic features, such as chromatin accessibility. Furthermore, when *O*^6^-MeG distributions in TMZ-exposed cells were normalized by comparing them to *O*^6^-MeG distributions in TMZ-exposed naked DNA of the same origin, *O*^6^-MeG seemed to be almost homogenously distributed throughout the genome. These observations suggest that *O*^6^-MeG formation is driven by intrinsic reactivity preferences of TMZ with DNA, and in particular certain trinucleotide contexts, rather than by the cell environment, and that MGMT removes *O*^6^-MeG without impacting its genomic distribution.

Furthermore, we found *O*^6^-MeG accumulated more in the non-transcribed strand in expressed genes, which also matches the strand bias found in SBS 11.(25) Likewise, other DNA adducts were found to be more abundant in the non-transcribed strand indicating involvement of transcription-associated repair like transcription-coupled nucleotide excision repair (TC-NER).(14,17,19,22,42) TC-NER is triggered by stalling of the RNA polymerase II during transcription, and indeed, it has been reported that human RNA polymerase II partially stalls at *O*^6^-MeG.(43,44) Hence, active TC-NER could be repairing *O*^6^-MeG in the transcribed strand leading to the strand bias we find in our data. While it has been shown that *O*^6^-carboxymethylguanine as well as AP sites can be repaired by NER (45,46), no such study has been conducted for *O*^6^-MeG. Further studies could investigate *O*^6^-MeG accumulation in cells lacking proteins of TC-NER. Finally, we found oncogenes to have less *O*^6^-MeG in LN-229 +MGMT than LN-229 WT (Figure 4J), suggesting that MGMT protects against mutations in oncogenes. If similar preferences arise in normal or stem cell populations, from which secondary cancers following TMZ exposure arise, MGMT could be a protective factor against carcinogenesis.

With a new method to map *O*^6^-MeG genome-wide and at single-nucleotide resolution, we analyzed the distribution of *O*^6^-MeG upon TMZ exposure in a glioblastoma cell line, and tested how its levels, signatures, and accumulation in particular genomic regions were impacted by expression of the *O*^6^-MeG repair enzyme MGMT. We could show that MGMT effectively reduces overall *O*^6^-MeG levels, while its impact on the distribution or sequence contexts of *O*^6^-MeG remains limited. Our data revealed distinct MGMT-independent trinucleotide preferences in *O*^6^-MeG formation in cells as well as in exposed naked DNA. Furthermore, a newly described *O*^6^-MeG signature strongly links the origin of SBS 11 to TMZ exposure. The identification of an MGMT-independent strand bias in *O*^6^-MeG accumulation within expressed genes suggests an additional transcription-associated repair mechanism for *O*^6^-MeG. However, reduced accumulation of *O*^6^-MeG in oncogenes in LN-229 +MGMT cells in comparison to LN-229 WT suggest a potential protective function of MGMT against mutations in oncogenes. Further application of *O*^6^-MeG-seq could help address additional factors in cancer or normal cells, such as how MMR or TC-NER proficiency and regulation of gene expression impacts drug resistance or avoidance of secondary cancers associated with chemotherapy use.

## DATA AVAILABILITY

*O*^6^-MeG-seq raw sequencing files and processed files can be found on NCBI Gene Expression Omnibus (GEO) (accession number GSE249155). All other data and support files can be found on Zenodo (DOI 10.5281/zenodo.10518966) with restricted access. Data analysis scripts and notebooks are also available at https://gitlab.ethz.ch/eth_toxlab/o6meg-seq

## AUTHOR CONTRIBUTIONS

J. Büchel and C. Mingard performed cell culture-based assays, prepared and analyzed samples by LC-MS/MS, did primer extension studies, prepared sequencing libraries, performed sequencing data analysis, interpreted data and wrote the manuscript. V. Takhaveev developed data analysis scripts, interpreted data and wrote the manuscript. P.B. Reinert, performed cell culture-based assays, prepared LC-MS/MS samples, prepared sequencing libraries and interpreted data. G. Keller performed primer extension studies and prepared sequencing libraries. T. Kloter performed primer extension studies and interpreted data. S. Huber developed the LC-MS/MS method. M. McKeague advised on experiment design. S. Sturla conceived, designed and supervised the study, interpreted data and wrote the manuscript.

## ACKNOWLEDGEMENTS

We thank Prof. Michael Weller (Laboratory of Molecular Neuro-Oncology, University Zurich) for providing the LN-229 cell lines. We acknowledge the Functional Genomics Center Zurich (FGCZ) and Genetic Diversity Centre (GDC), sequencing and experimental instrumentation platforms used for this research, and their staff for technical support.

