## Supplementary_figures for "Single-nucleotide-resolution genomic maps of *O*^6^-methylguanine from the glioblastoma drug temozolomide"

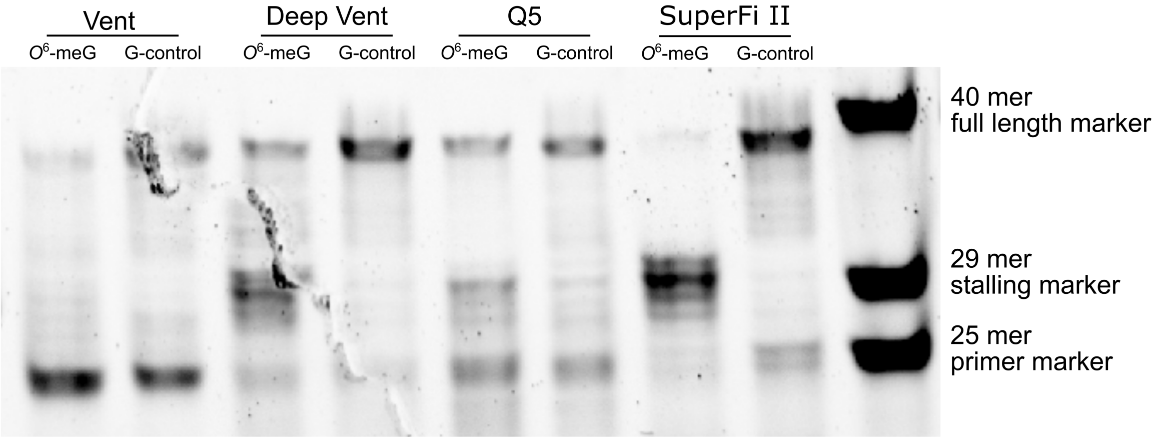


**Supp. Figure 1** Polymerase extension assay**.** Vent, Deep Vent, Q5 and SuperFi II polymerase were used for primer extension on a 40mer template containing either *O*$^6$-MeG or a normal guanine (G-control). Only SuperFi II was stalled at *O*$^6$-MeG while the G-control was extended in full length.

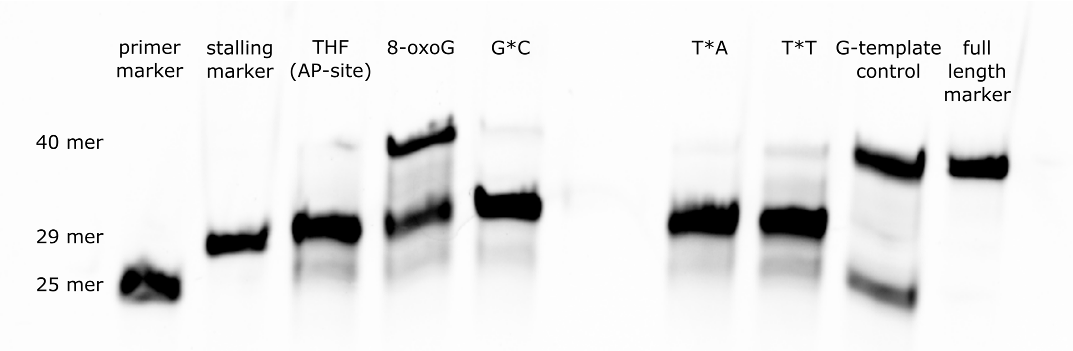


**Supp. Figure 2** SuperFi II polymerase stalling at different modifications and contexts. 40mer templates containing tetrahydrofuran (THF, resembling an abasic (AP) site), 8-oxoguanine (8-oxoG) and *O*$^6$-MeG (indicated as asterisk) in different trinucleotide contexts together with the G-template control were tested for SuperFi II polymerase stalling.

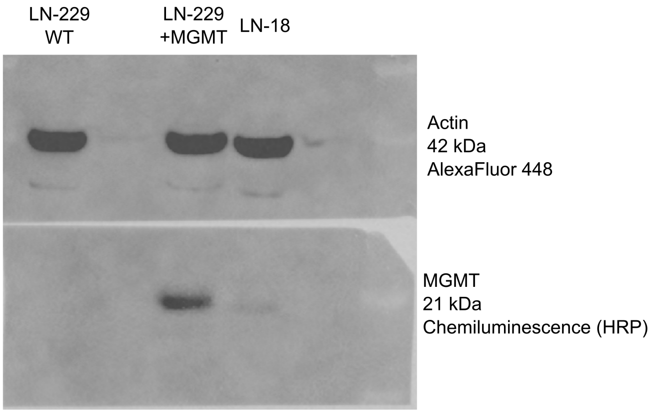


**Supp. Figure 3** Western blot for MGMT protein. Whole cell extracts of LN-229 WT and +MGMT cells, as well as LN-18 with known MGMT expression, were analyzed by western blot to confirm MGMT expression.

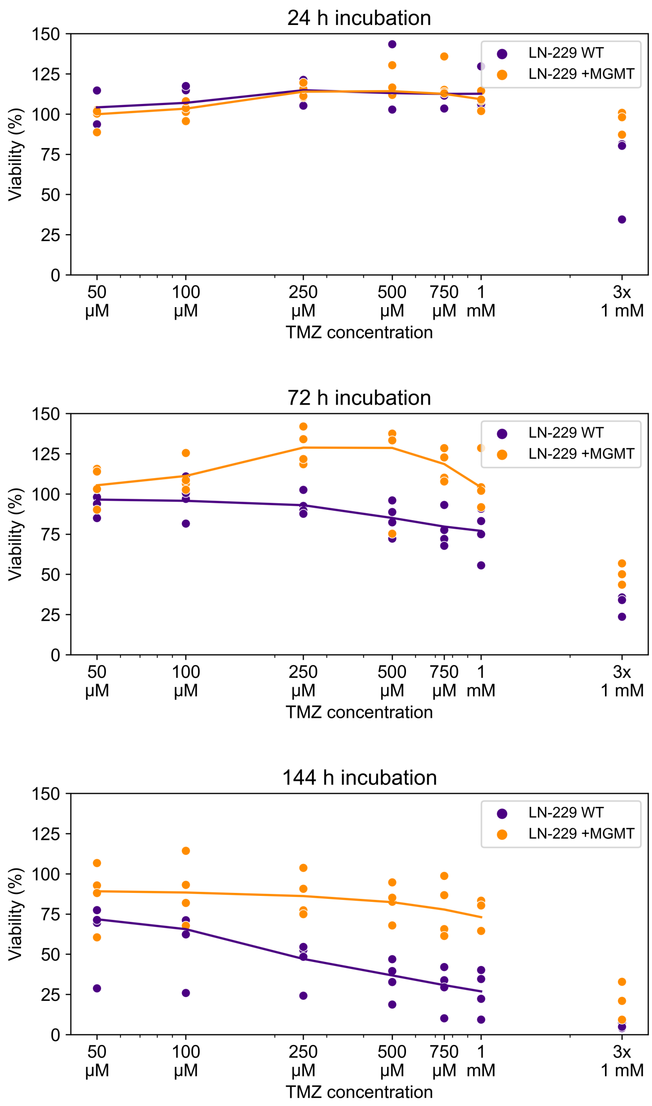


**Supp. Figure 4** LN-229 cell viability. LN-229 cells were exposed to TMZ in the medium and incubated for 24 to 144 hours. 4 replicates for all TMZ concentrations except 3x 1mM where n=3. Cell viability was assessed with CellTiter-Glo measuring relative ATP concentrations.

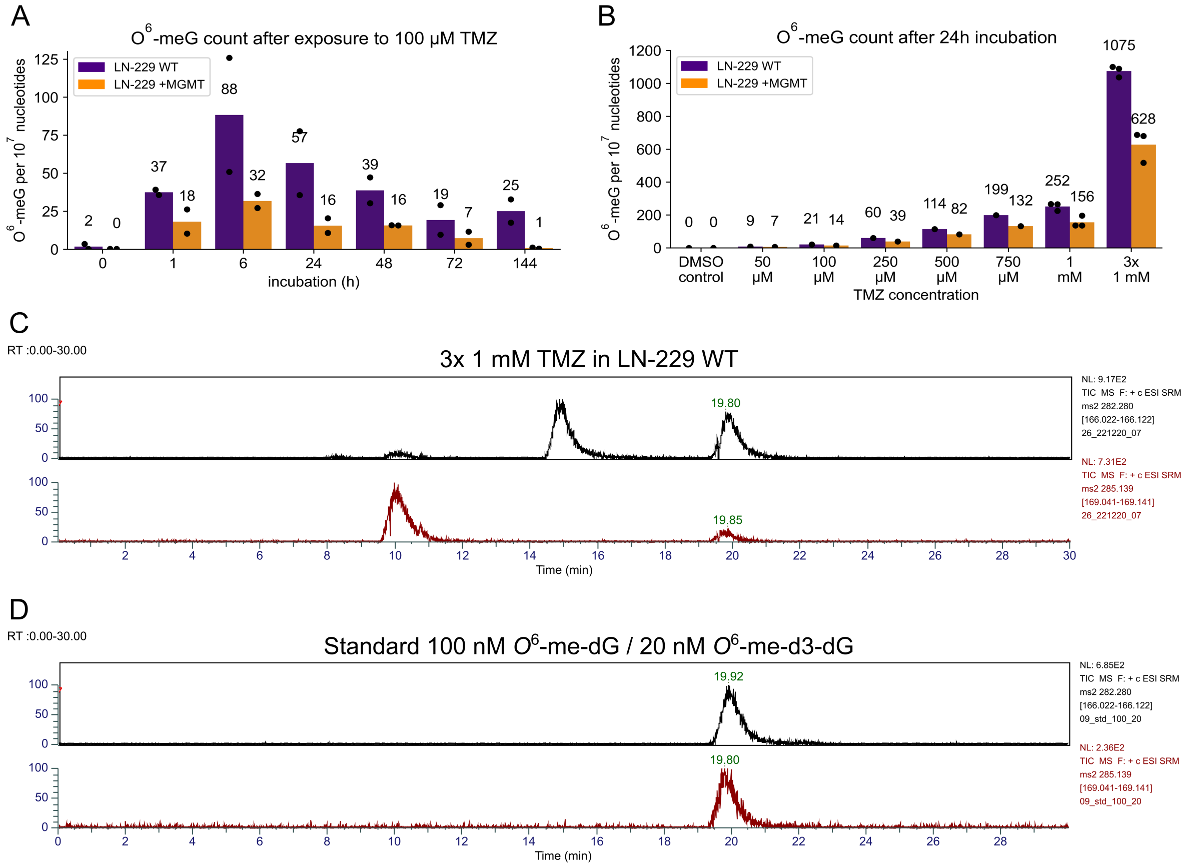


**Supp. Figure 5** *O*$^6$-MeG quantification with mass-spectrometry. **A** *O*$^6$-MeG counts after exposure to 100uM TMZ for 0 – 144 h. **B** *O*$^6$-MeG counts after exposure to various TMZ concentrations for 24 h. **C** *O*$^6$-MeG counts of naked DNA exposed to 1 mM TMZ for 24h. **D** Representative chromatograms of *O*$^6$-MeG quantification in 3x 1 mM TMZ- exposed LN-229 WT. Top ion chromatogram (black): single reaction monitoring (SRM) m/z 282 to 166 for *O*$^6$-me-dG; bottom ion chromatogram (red): SRM m/z 285 to 169 for *O*$^6$-me-d3-dG as internal standard. **E** Representative chromatogram of standard used for *O*$^6$-MeG quantification including 100 nM *O*$^6$-me-dG (top chromatogram in black) and 20 nM *O*$^6$-me-d3-dG (top chromatogram in red).

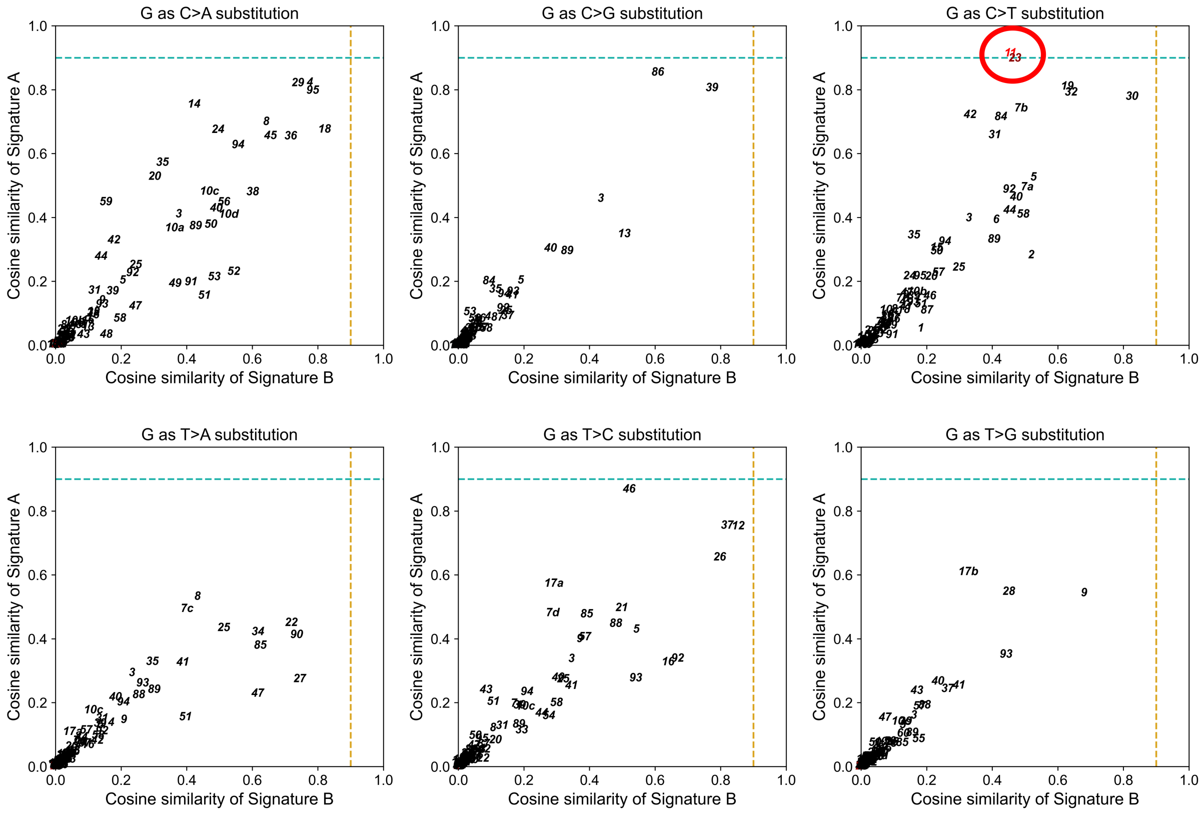


**Supp. Figure 6** Cosine similarities of $O^6$-MeG signatures A and B compared to COSMIC SBS signatures. As the signatures cannot be directly compared due to their different dimensions, the XGY contexts of the $O^6$-MeG signatures were converted into X′[C>T]Y′, X′[C>A]Y′, X′[C>G]Y′, X′[T>A]Y′, X′[T>C]Y′ or X′[T>G]Y′ mutations, where X′ and Y′ are reverse complements of the flanking bases present in the modified triplets. All other contexts were set to zero. Only the conversion to X′[C>T]Y′ revealed high similarities with COSMIC SBS 11 and 23 (red circle). Cosine similarity of 0.9 was used as cut off for high similarity (dashed lines).

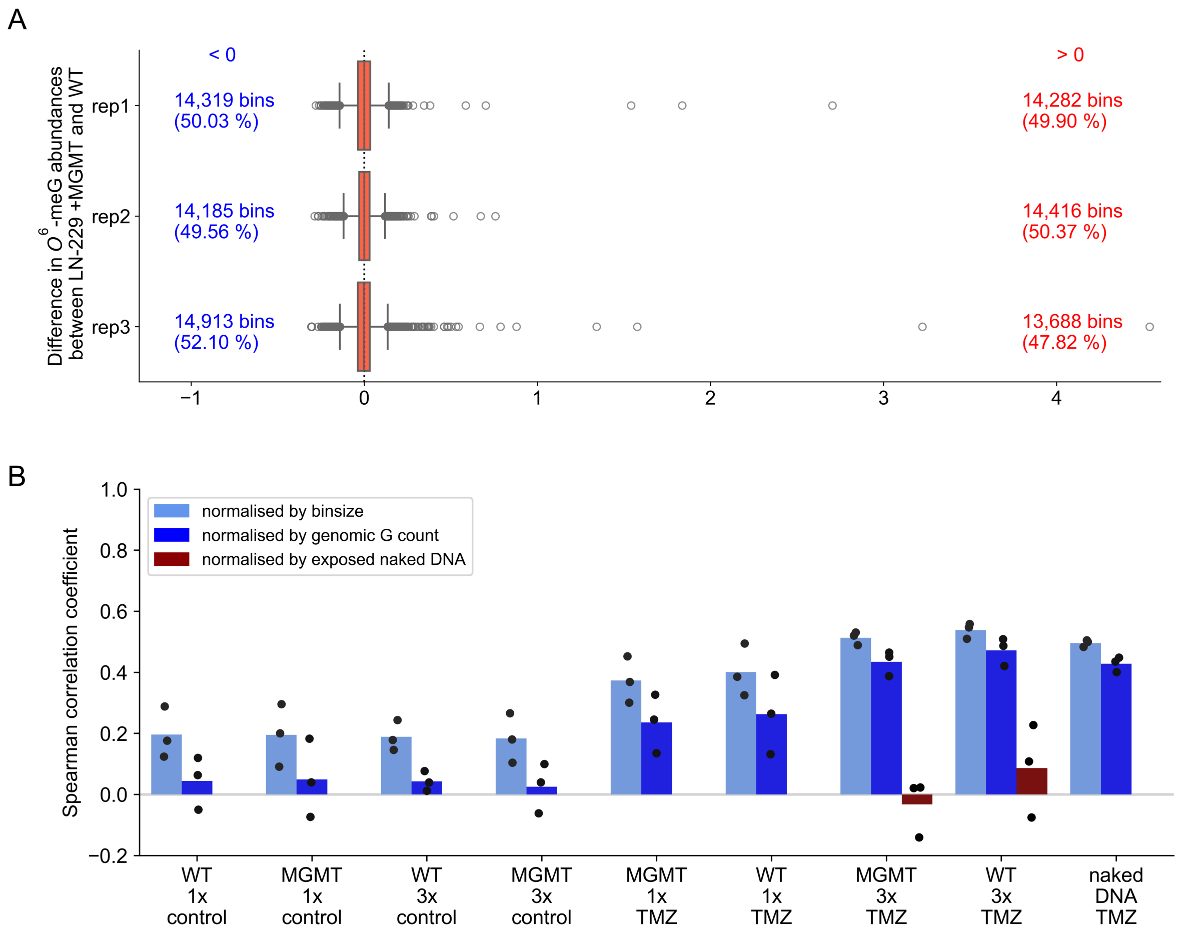


**Supp. Figure 7** *O*$^6$-MeG abundance in 100 Kb bins in comparison. **A** *O*$^6$-MeG-seq data from LN-229 +MGMT cells compared to LN-229 WT cells by subtracting *O*$^6$-MeG abundance in 100 Kb bins normalized by TMZ-exposed naked DNA. **B** Spearman correlation of *O*$^6$-MeG abundance (normalization methods as in Figure 3) with ATAC-Seq data in 100 Kb bins. P values $<10^{-3}$ for all comparisons.

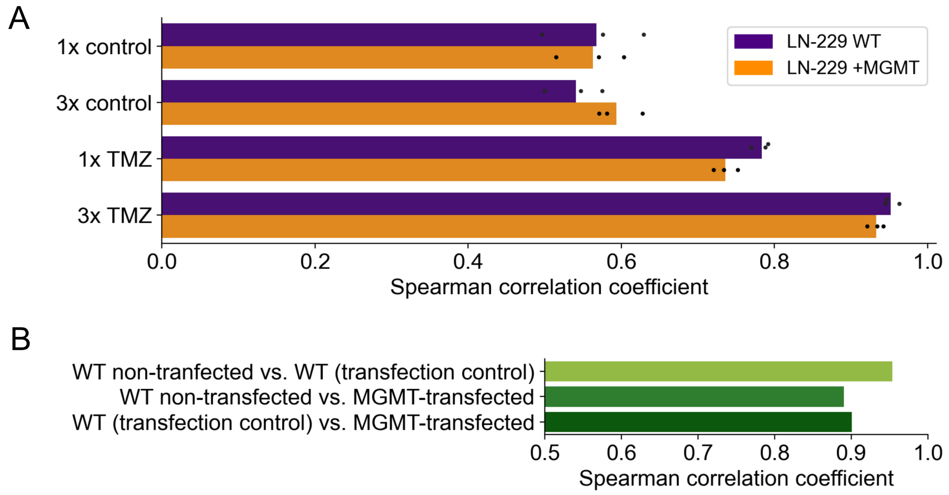


**Supp. Figure 8** Spearman correlation of replicates. P values $<10^{-5}$ for all comparisons. **A** Spearman correlation coefficients of cell sample replicates. **B** Spearman correlation coefficients of exposed naked DNA of different cell origin.
